# Negative feedback on Retinoic Acid by Brachyury guides gastruloid symmetry-breaking

**DOI:** 10.1101/2023.06.02.543388

**Authors:** Meagan J. Hennessy, Timothy Fulton, David A. Turner, Ben Steventon

## Abstract

Establishment of the vertebrate body plan requires a combination of extra-embryonic signalling to establish morphogen gradients, and an underlying self-assembly mechanism that contributes to pattern regulation and robustness. Gastruloids are aggregates of mouse embryonic stem cells that break morphological symmetry and polarise *Brachyury* (*Bra*) expression in the absence of extra-embryonic signals. However, the mechanism by which symmetry breaking occurs is not yet known. During gastrulation and body axis elongation, retinoic acid (RA) and *Cyp26a1* are polarised along the anteroposterior axis, and this is critical for balancing the decision of cells to self-renew or differentiate. We found that symmetry-breaking in gastruloids is coincident with the separation of *Aldh1a2* and *Cyp26a1* expression, and that feedback from *Bra* is critical for maintaining polarised *Cyp26a1* gene expression in the gastruloid posterior region. Furthermore, we reveal a short temporal window where RA signalling can negatively influence both *Bra* and *Cyp26a1* expression. These observations lead us to suggest a mechanism of how initial gastruloid patterning, subsequent elongation, and evolving network topologies can create defined boundaries of RA signalling that permits proper axial patterning and gastruloid growth.

## 1. Introduction

Development of the body plan requires the integration of morphogen gradients and cell fate decision-making together with large scale tissue deformations to generate multi-tissue morphogenesis. This interplay enables a large degree of regulative capacity that generates both robustness and evolvability (Steventon et al., 2021). Recently, this has been further exemplified in the development of stem cell protocols that enable the self-assembly of rudimentary multi-axial patterning and subsequent organogenesis (Moris et al., 2020; Veenvliet et al., 2021). Gastruloids are 3D aggregates of mammalian embryonic stem cells (ESCs) that can break morphological symmetry and polarise early mesodermal markers in the absence of extraembryonic signals (Beccari et al., 2018; Turner et al., 2017; van den Brink et al., 2014). Recent work has shown how the balance between pluripotency and differentiation in early gastruloids can illicit a bimodal response to signals to aid their symmetry breaking (Suppinger et al., 2023). In addition, differential cell adhesion has been shown to play a key role in organising cell movements during the process (Hashmi, et al. 2022). However, the ways in which multi-cellular communication enable self-assembly in this context are not yet fully known. Uncovering such mechanisms will shed light on the regulative mechanisms of body plan formation during normal development.

A defined set of signalling pathways are utilised by the embryo during early development which orchestrate axial patterning, morphogenesis and the subsequent unfolding of the body plan. Retinoic acid (RA) is a key molecule that provides signalling inputs at multiple stages of development, controlling axial patterning, morphogenesis, and cell fate (Duester, 2008). It is critical for the embryo to balance the temporal and spatial distribution of RA synthesis as experimental perturbations which target RA (either positively or negatively) cause wide-ranging developmental abnormalities. Such control of RA is brought about through the activity of enzymes controlling its synthesis (e.g. aldehyde dehydrogenases; Aldh) and its degradation (cytochrome P450; Cyp) which are expressed in a temporal and tissue-specific manner. In the mouse embryo, RA activity is not detected prior to embryonic day (E) *∼* 7.5, however following the formation of the mesoderm, its activity can be found along the primitive streak, and throughout the posterior portion of the embryo and the nascent mesoderm (Rossant et al., 1991). At later stages of development, RA activity is reduced in the tailbud, expressed in all tissues of the embryo trunk up to the boundary between the hind-brain and the first somite, and within optic lobes and surrounding mesenchyme in the prospective head region (Rossant et al., 1991). This pattern of expression is mirrored by the presence of the enzymes controlling its synthesis and degradation. Prior to gastrulation, *Cyp26a1* is expressed throughout the whole embryo as a protective barrier against maternal RA (Uehara et al., 2009). By E7.25 *Cyp26a1* expression can be seen in the primitive streak following its progression anteriorly by E7.5 (Fujii et al., 1997), and by E9.0, it’s confined to the tailbud (Sakai et al., 2001). *Aldh1a2* on the other hand is first detected in the primitive streak and nascent mesoderm of E7.5 embryos, co-localises with RA signalling, forming a distinct signalling boundary with the expression of *Cyp26a1* (Ribes et al., 2009). Taken together with loss/gain of function studies (Iulianella et al., 1999; Molotkova et al., 2005; Rossant et al., 1991; Sakai et al., 2001), it is clear that the timely positioning of metabolic enzymes controlling its synthesis and activity is functionally important for enabling proper anteroposterior patterning. However, the molecular mechanisms that bring about both the separation and maintenance of *Aldh1a2* and *Cyp26a1* expression domains prior to, and during axial elongation are not fully understood.

Given the known importance of establishing a polarity in RA production during mouse gastrulation, we sought to understand the impact of RA signalling on symmetry breaking in gastruloids and test the hypothesis that feedback from T/Brachyury (Bra) to the control of retinoic acid production might play an important role in symmetry breaking and early mesoderm specification.

## 2 Results and Discussion

### 2.1 Progressive formation of spatial boundaries between Aldh1a2 and Cyp26a1 in gastruloids

To gain a better understanding of the regulation of *Aldh1a2* and *Cyp26a1* expression in gastruloids as they undergo symmetry breaking, we characterised their expression in gastruloids over time (**Fig. 1**), choosing time-points that mapped to stages of the early embryo that spanned the initiation of gastrulation (72h *∼* E6.5) to axial patterning (120h *∼* E8.5) (Martinez Arias et al., 2022). We took a fluorescence-based *in situ* Hybridisation Chain Reaction (HCR) approach to detect multiple transcripts within samples (Choi et al., 2018; Fulton et al., 2020).

**Figure 1:**
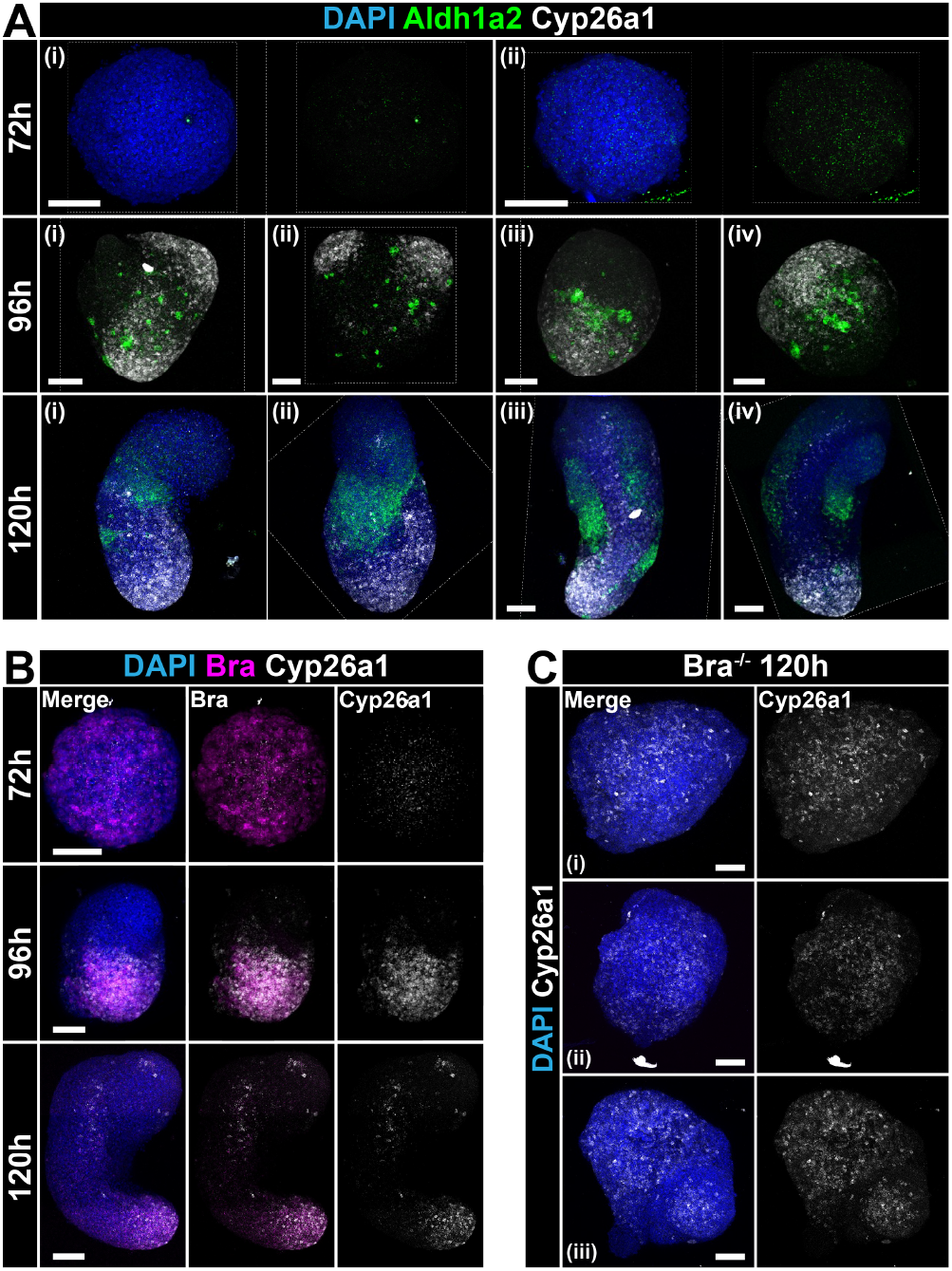
*In situ* hybridisation chain reaction (HCR) analysis reveals the temporal Progression of *Aldh1a2* and *Cyp26a1* expression. Gastruloids made from wild-type E14Tg2A mouse ESCs were fixed at the indicated times after aggregation (AA) and probed for *Cyp26a1* with either (**A**) *Aldha2* or (**B**) *Brachyury* (Bra). There is a polarisation of *Cyp26a1* which overlaps with *Bra* expression, and a clear boundary of expression between *Aldha2* and *Cyp26a1*. Two or four representative images shown for each time-point in (A), indicated by numerals. (**C**) Gastruloids at 120h AA formed from *Bra*^-/-^ mouse ESCs fail to elongate at 120h, and express non-polarised and low levels of *Cyp26a1*. Three representative gastruloids shown; scale bar indicates 100μm in all images; nuclei stained with DAPI (blue).

At 72h after aggregation (AA), gastruloids showed a typical spherical morphology, with scattered, low-level expression of both *Aldh1a2* and *Cyp26a1* throughout the gastruloid, with little evidence of polarisation or distinct expression pattern domains (**Fig. 1A**; top row). To orientate the expression of these markers to either ‘anterior’ or ‘posterior’ regions of gastruloids, we also included probes for *Bra*, a marker of the primitive streak and nascent mesoderm (Wilkinson et al., 1990), and by definition, a land-mark for the posterior of gastruloids (van den Brink et al., 2014) (**Fig. 1B**). *Bra* was expressed throughout gastruloids by 72h AA, with only a marginal indication of asymmetry in its expression, and there appeared little correlation between its localisation and *Cyp26a1* or *Aldh1a2* (**Fig. 1B; fig. S1B**). Gastruloids had initiated elongation by 96h AA, and there was strong up-regulation of *Cyp26a1* and clear signs of polarisation in multiple gastruloids. By contrast, *Aldh1a2*, although had higher expression than at 72h AA, it was expressed in small clusters of cells anterior to the *Cyp26a1* domain (**Fig. 1A**; middle row), mapping to regions of gastruloids with lower expression of *Cyp26a1* (**Fig. 1A**; middle row). By 120h AA, gastruloids had their characteristic elongated morphology, and showed expression domains of both *Cyp26a1* and *Aldh1a2* (**Fig. 1A**; bottom row; **fig. S1A**), with the former confined to Bra expression domain in the elongating posterior region (**Fig. 1B**; middle row), and the latter in distinct, non-overlapping regions towards the more anterior, trunk sections of the gastruloid.

Taken together, these data indicate that the up-regulation and spatial separation of *Aldh1a2* and *Cyp26a1* is co-incident with symmetry-breaking (indicated by *Bra* polarisation) and axial elongation.

### 2.2 Brachyury is necessary for Cyp26a1 polarisation

The spatially overlapping, polarised co-expression of *Cyp26a1* with *Bra* led us to hypothesise that *Bra* may be driving or maintaining the expression of *Cyp26a1* in the posterior region of gastruloids, creating a region of low RA signalling similar to what has been reported in Zebrafish (Martin and Kimelman, 2010). To examine this, we generated gastruloids from mouse ESCs isolated from *Bra*^-/-^ embryos (Rashbass et al., 1991; Wilkinson et al., 1990), cultured them in standard gastruloid conditions for 120h AA, and probed them for *Cyp26a1* using HCR (**Fig. 1C**). In agreement with previous observations (Wehmeyer et al., 2022), the absence of *Bra* resulted in a complete inhibition of gastruloid elongation. This may reflect the phenotype of *Bra*^-/-^ embryos where Bra protein coordinates the movement of cells away from the primitive streak during gastrulation (Beddington et al., 1992; Wilson and Beddington, 1997; Wilson et al., 1995; Yanagisawa et al., 1981), and in primitive streak differentiation assays in culture (Hashimoto et al., 1987; Turner et al., 2014b). Although *Cyp26a1* was still expressed in Bra^-/-^ gastruloids (albeit at low levels of detection), there was no indication of polarisation (**Fig. 1C**).

Next, to examine whether this co-dependency can be seen in an environment where coordinated polarisation is minimal (e.g. 2D cultures), we cultured wild-type E14-Tg2A mouse ESCs as monolayer, and exposed them to continuous 3µM Chi, a Wnt/β-Catenin agonist, which is known to induce the expression of *Bra* (Turner et al., 2014a; Turner et al., 2014b; **fig. S1C**). Whereas *Bra* and *Cyp26a1* were up-regulated in clusters within ESC colonies, however *Cyp26a1* was also expressed in colonies where *Bra* was absent (**fig. S1C**).

Taken together, these data suggest that although *Cyp26a1* is not dependent on *Bra* for its initial expression in gastruloids, *Bra* is necessary for its polarisation and enhanced transcription in 3D, in agreement with previous studies (Martin and Kimelman, 2010), which may constitute a feedback loop to reduce RA expression.

### 2.3 Gastruloids Exhibit Temporal Sensitivity to RA Addition

As symmetry-breaking in gastruloids is coincident with the polarisation of *Cyp26a1* and *Bra*, we next asked whether RA signalling may modulate the expression of *Bra*. It is known that the levels of RA must be controlled both spatially and temporally for normal development, and exposure to excess RA during embryogenesis gives rise to different phenotypes depending on the time of exposure and the concentration used (Huang et al. 2001). Mouse embryos that are null mutants of RARγ also display a reduction in *Bra* expression (Iulianella et al., 1999). Furthermore, RA is a potent teratogenic agent, and a recent study assessing the effect of teratogenic agents on gastruloids using a range of RA doses showed that high RA disrupts axial extension and negatively regulated the expression of posterior markers (*Bra*) in favour of neural (*Sox2*) (Mantziou et al., 2021). Indeed, these observations also hold after we treated gastruloids with low concentrations of RA from 72-120 AA, showing a loss axial elongation (**fig. S2**).

To determine whether there were key time intervals where gastruloids were most sensitive to RA exposure in terms of their elongation and polarisation potential, we exposed gastruloids to single 24h pulses of low-dose RA between 72-96h or 96-120h AA (**Fig. 2A-D**). Early exposure to RA between 72-96h AA disrupted gastruloid elongation as well as the expression and polarisation of *Bra* and *Cyp26a1*, whereas later treatment (96-120h AA) resulted in more elongated gastruloids (**Fig. 2D**) and significantly higher level of *Cyp26a1* expression in the posterior region (**Fig. 2A,C**). Interestingly, although the induction of *Bra* was elevated following treatment of RA between 96-120h AA, compared with 72-120h or 72-96h AA, *Bra* levels were significantly lower than controls, suggesting that inhibition or a reduction RA are required for *Bra* maintenance in the tailbud region (**Fig. 2A,D**). We reasoned that the elevated levels of *Cyp26a1* in gastruloids exposed to RA between 96-120h AA may be accounted for by the dual action of positive regulation on *Cyp26a1* by *Bra* and RA (Martin and Kimelman, 2010). The differential response to RA may in part be due to the initial patterning of gastruloids at the time of RA induction: early inhibition of *Cyp26a1* prevents *Bra* up-regulation, negatively influence axial extension in the embryo (Beddington et al., 1992; Wilson and Beddington, 1997; Wilson et al., 1995). This is also reflected by truncated gastruloids formed from *Bra*^-/-^ ESCs (**Fig. 1c**).

**Figure 2:**
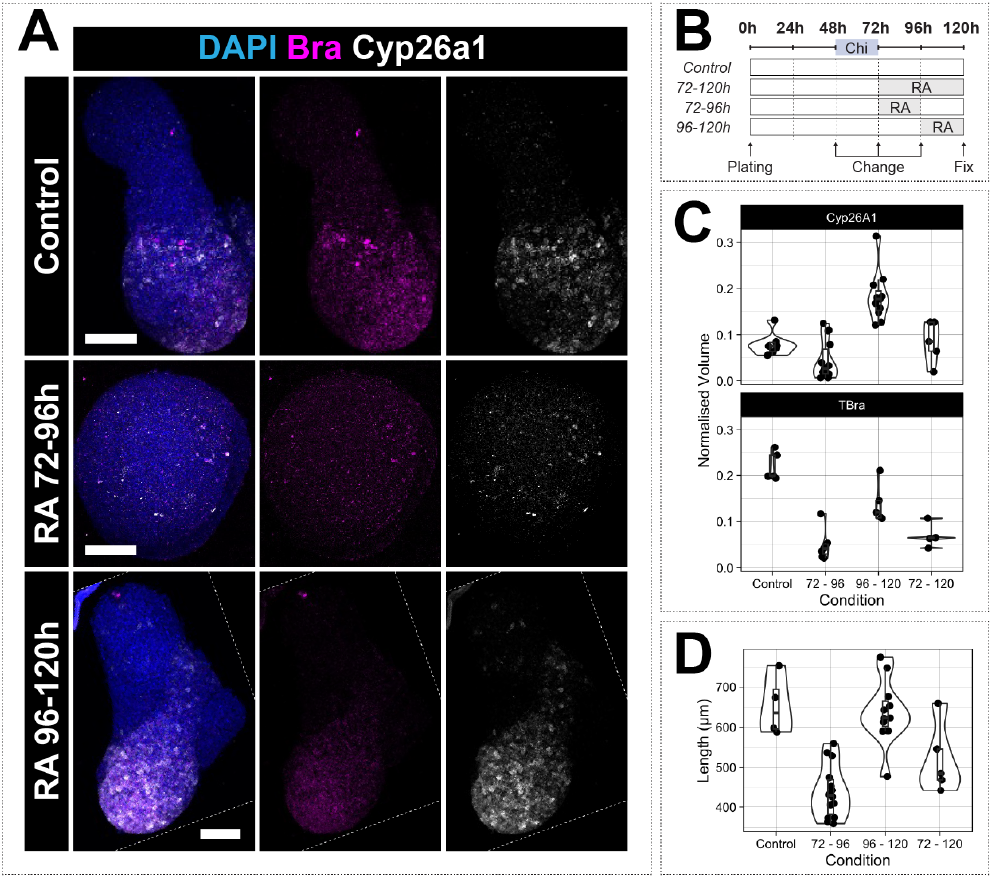
Gastruloids display temporal sensitivity to low-dose Retinoic Acid. Gastruloids fixed at 120h AA, probed for *Bra* and *Cyp26a1* (**A**) were treated as indicated (**B**) with RA at different intervals. The normalised volume of *Cyp26a1* (**C**, top) and *Bra* (**C**, bottom) were calculated, as well as the gastruloid length (**D**). Scale bars indicate 100μm.

In summary, our observations support the notion that there is a temporal window where RA can negatively regulate both *Cyp26a1* and *Bra* expression which maps to the symmetry breaking step of gastruloid development.

### 2.4 RA inhibition disrupts gastruloid volume and Cyp26a1 expression, but not elongation

Our approach so far has been to add excess signalling at defined temporal windows, however it is not yet clear how gastruloids would respond to a RA-free environment, and whether basal, endogenous (low) levels of RA can have a positive influence on gastruloid symmetry-breaking.

We therefore exposed gastruloids the potent pan-RA receptor antagonist AGN 193109 (AGN; Johnson et al., 1995; Soprano et al., 2001) tonically for 72h (between 48-120h AA) or as a single 24h pulse on subsequent days (48-72h, 72-96, 96-120h AA; **Fig. 3**). Treatment with AGN between 48-120h AA resulted in gastruloids that were unable to maintain the level of *Cyp26a1* or *Bra*. AGN pulsed between 48-72h, 72h-96 and 96-120h AA significantly reduced the levels and expression domains of *Cyp26a1* (**Fig. 3A,B**). *Bra* expression on the other hand, although reduced following a 48-72h pulse of AGN, were generally not greatly influenced by loss of RA signalling, consistent with previous observations that *Bra* is a target of RA-induced negative regulation (Iulianella et al., 1999). We also observed a loss of co-localisation between *Bra* and *Cyp26a1* if RA is inhibited between 48-72 or 72-96 AA. Finally, loss of RA signalling appeared to have a negative influence on the size of gastruloids, although their potential to elongate did not appear to be greatly disrupted (**Fig. 3**). Taken together, our data suggest that the expression of *Cyp26a1* and its co-localisation with Bra may benefit from low levels of RA signalling.

**Figure 3:**
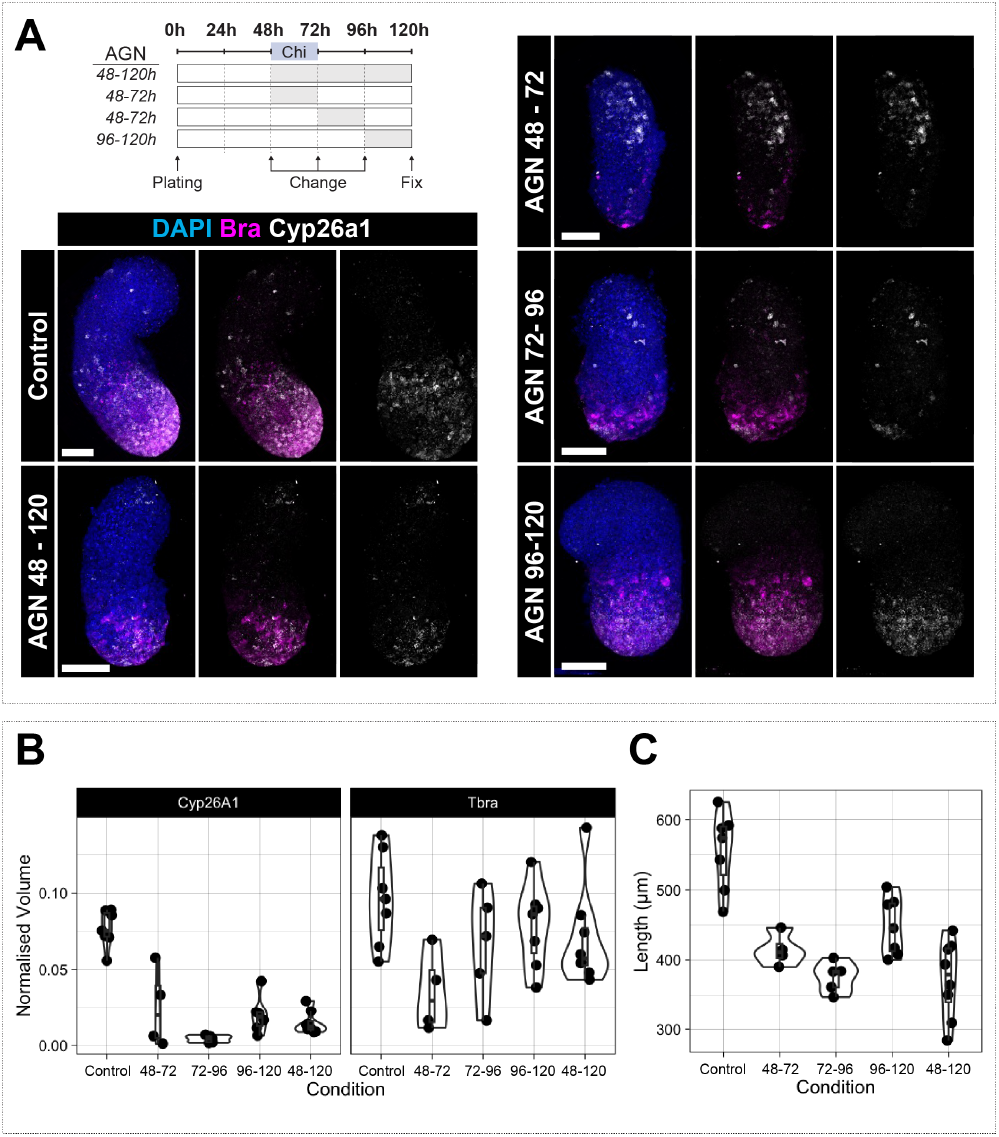
Low-dose RA signalling is required for growth and maintenance of gene expression. Gastruloids fixed at 120h AA and probed for *Bra* and *Cyp26a1* (**A**) were treated as indicated with AGN at different intervals and the normalised volume of *Cyp26a1* (**B**, left) and *Bra* (**B**, right) were calculated. Long-term inhibition of RA reduces expression of *Bra* and *Cyp26a1* and length (**C**). Gastruloids most sensitive to RA inhibition between 48-72h AA.

## 3. Conclusions and perspectives

We have used mouse gastruloids to probe the mechanisms through which RA signalling may be regulated during (and also influence) early axial development. We find that:

1. Separation of *Aldh2a1* and *Cyp26a1* is co-incident with symmetry breaking in gastruloids;
2. *Bra* is required for *Cyp26a1* polarization, but not its initial expression;
3. RA has a temporal window between 48 and 96h where it can negatively influence both *Cyp26a1* and *Bra* expression;
4. Prolonged RA inhibition results in a decrease in gastruloid size and Cyp26a1 expression, with marginal effect on *Bra*, and inhibition between 72-96h disrupts *Cyp26a1* polarization.

From these observations, we propose a model which includes a simple gene regulatory network (GRN) describing how *Aldh1a2* and *Cyp26a1* maintain RA signalling. In the first instance, *Aldh1a2* and *Cyp26a1* are heterogeneous, with low expression, and probably in equilibrium, providing low-level endogenous RA signalling across the gastruloid (**Fig. 4**; left). Inductive signals between 48 and 72h from Wnt/β-Catenin (Chi pulse) activates *Bra* which feeds into the base GRN module, initially elevating *Cyp26a1* in regions where *Bra* expression is highest leading to asymmetry in *Cyp26a1* expression, and therefore in RA levels (**Fig. 4**; middle). It is assumed there is a further GRN module, a Wnt3a-Bra autoregulatory loop, independent of the *Aldh1a2*-RA-*Cyp26a1* network that maintains *Bra* expression in the elongating tailbud (Yamaguchi et al., 1999; Martin and Kimelman, 2008), but is not included here for clarity. The continued presence of *Bra*, however, drives gastrulation-like processes and facilitates axial elongation (**Fig. 4**; right).

**Figure 4:**
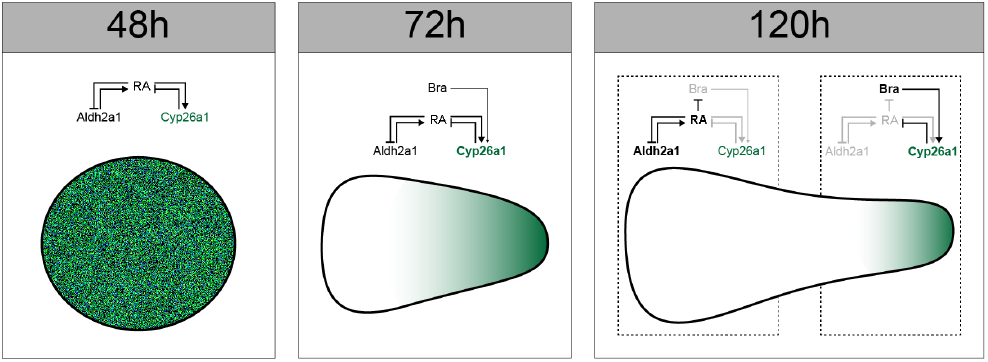
Suggested model to describe segregation of *Cyp26a1* and *Aldh1a2* during axial elongation and patterning in gastruloids. Possible network topology showing positive and negative feedback regulating RA (Left panel, 48h). By 48h AA, there is low, heterogeneous expression of *Cyp26a1* (green) and *Aldh1a2* (non-green). At the end of the Chi-pulse (at 72h), *Bra* is induced and is an additional input into the original network topology, increasing the effect *Cyp26a1* has in the network (indicated by bold font) which starts to create asymmetries between *Aldh1a2* and *Cyp26a1*. As gastruloids elongate, positive feedback within the posterior region maintains Bra expression and RA inhibition via *Cyp26a1*. Lower *Bra* signalling in the anterior region (by virtue of being further away from elongating tip) tilts the balance of the network in favour of *Ald1a2* expression and increased RA signalling. *Bra* may therefore serve as a mediator in establishing the boundaries between *Cyp26a1* and *Aldh1a2* domains, and thereby the concentration of RA available to cells during axial development. Bold text and bold arrows indicate a stronger input into the network; greyed out text and arrows suggest minimal involvement in the network.

As elongation progresses and the *Bra*/*Cyp26a1* domain is further away from the anterior region, a minimum threshold of Bra is reached causing in a switch in the GRN network topology where RA has more of an influence in suppressing Bra, leading to enhanced *Aldh1a2* expression and a clear boundary of expression domains. This mimics what is known about the regulation of RA gradient formation during axis elongation in zebrafish embryos and supports the notion that axial elongation is a important regulator of patterning through controlling the spatial segregation of signalling domains across the anteriorposterior axis (White et al., 2007). Indeed, blocking elongation of whole-embryo explants from zebrafish embryos has been *BioRxiv Submission* 5 shown to impact the patterning of neural marker expression (Fulton et al., 2020). Together, these findings lend support to the idea that multi-tissue morphogenesis, while initiated by early signalling events, can feed-back to the spatial and temporal regulation of signalling pathways to provide a mechanism of pattern regulation through multi-tissue tectonics (Busby and Steventon, 2021).

In summary, by using a gastruloid model, we have demonstrated the temporal requirements for RA signalling and how it relates to axial patterning and cell fate. We see this as one of likely many contributing mechanisms that enable symmetry breaking in gastruloids. As such, our proposed model can serve as a baseline for further studies to uncover the dynamics of these patterning processes and GRNs, which will allow a more detailed understanding of how growth, morphogenesis, metabolic control of signalling are coupled to determine the self-assembly of the embryonic body plan.

## 4. Materials and Methods

### 4.1 Routine mouse embryonic stem cell culture

Wild-type E14-Tg2A (Hooper et al., 1987) and BR BT10 (Bra^-/-^) (Beddington et al., 1992; Rashbass et al., 1991) mouse ESCs were seeded at a density of 1.2x10^4^ cells/cm^2^ on 0.1% (v/v) gelatin-coated flasks in ESL medium. ESL contains a base medium of GMEM (Gibco, 21710082) supplemented with 10% Foetal Bovine Serum (BioSera, FB-1090/500), Non-Essential Amino Acids (Gibco, 11140035), Sodium Pyruvate (Gibco, 11360039), Glutamax™ (Gibco, 35050038), 2 mercaptoethanol (Gibco, 31350010), and LIF (QKine; QK018) in a humidified incubator maintained at 37°C with 5% CO_2_. Cells were passaged every second day, with full medium changes in between passages. Cells were tested monthly and certified negative for mycoplasma.

### 4.2 Generation of Gastruloids

Gastruloids were generated as previously described (Baillie-Johnson et al., 2015; Beccari et al., 2018; Turner et al., 2017; van den Brink et al., 2014) with the following modifications. Cells were sub-cultured from a stock flask and plated in ESL and 24h later, medium replaced with 2iLIF which contains 3µM CHIR 99021 (Chi; Tocris, 4423), 1µM PD 0325901 (PD03; Tocris, 4192), and LIF in a base medium of NDiff227 (Takara, Y40002) (Ying et al., 2008). The next day, culture medium was aspirated, and cells rinsed with phosphate buffered saline (PBS) without Mg^2+^ or Ca^2+^ (PBS^-/-^), detached using 2mL trypsin, and resuspended in 8mL ESL prior to centrifugation (5 min, 150g). Two successive rounds of aspiration, and resuspension in 10mL PBS^+/+^ and centrifugation followed to ensure all traces of growth medium was removed, prior to cells being fully resuspended to a single-cell suspension in 2mL NDiff227 with a 1mL pipette (Takara). Using a multichannel pipette, 40µL of a subsequent cell suspension was added to each well of a non-tissue culture treated 96-well plate, ensuring 350 cells/well, and left for 48h to permit aggregation. At 48h after aggregation (AA), all conditions received 150µL of secondary medium consisting of N2B27 supplemented with Chi (3µM) with or without other factors (see below). Secondary medium (150µL) was removed 24h later (72h AA), replaced with 150µL NDiff227, ensuring the gastruloid was agitated to prevent it sticking to the wells of the plate. Medium was replaced every day until 120h AA. When required, medium was supplemented with Retinoic Acid (RA; 0.4nM; Sigma-Aldrich, R2625) or AGN 193109 (AGN; 200nM; Sigma-Aldrich, SML2034).

### 4.3 In situ Hybridisation Chain Reaction (HCR)

HCR probes for *T/Brachyury* (*Bra*; **Table S1**) *Aldh1a1* (**Table S2**) and *Cyp26a1 (****Table S3***) were designed according to the parameters described in (Choi et al., 2018) with at ten split initiator pairs per transcript. Sequences were manually screened to ensure they were non-overlapping and spread across each gene’s transcript. The standard ‘version 3’ HCR protocol was followed according to those previously published (Choi et al., 2018) with minor modifications as described previously (Fulton et al., 2020). Briefly, samples were fixed in 4% (v/v) formaldehyde in PBS^-/-^ at room temperature in 500µL microcentrifuge tubes for 1h and incubated with 2pmol of probe over night at 37°C 500µL hybridization buffer (30% formamide), followed by repeated washes at 37C in probe wash buffer (30% formamide). Fluorescent hairpins targeting the probes were annealed in amplification buffer at room temperature, protected from light, over-night, followed by subsequent washes in 5x SSCT. Nuclei were stained with DAPI, and samples prepared for microscopy as described below. Fluorescent hairpins were purchased from Molecular Technologies.

### 4.4 Sample clearing, Confocal microscopy, Image Analysis

Samples were cleared over-night using ScaleS3 (Hama et al., 2015), and mounted in clearing reagent in #1.5 cover glass bottomed Petri dish as described previously (Fulton et al., 2020). Imaging was performed using an inverted LSM700 (Zeiss) confocal microscope on a Zeiss Axiovert 200M using a Plan-Apochromat 20x 0.8 NA air objective. DAPI and fluorescent hairpins conjugated to alexa-conjugated fluorophores -488 (B5), -546 (B2), and -647 (B4) were sequentially excited with 405, 488, 555 and 639-nm lasers, respectively, as previously described (Turner et al., 2017). Data were captured using Zen2010 and processed in the image analysis package Imaris (Bitplane).

The effect sizes between different treatments were estimated to show the variability in the response of different treatment regimes. The normalised volumes of gene expression (*Cyp36a1* and *Bra*) or gastruloid length were compared with the control plots for each treatment condition using a bootstrap-coupled estimation “dabestr” R package, and data visualised using Cumming estimation plots (Ho et al., 2019).

## Acknowledgement

We would like to thank Prof. Valerie Wilson for the kind gift of the Brachyury (BT BR10) mouse ESCs, Tina Balayo for expert advice in mouse ESC gastruloid culture, and Chaitanya Dingare for assistance in HCR probe design. We are grateful to Alfonso Martinez Arias useful discussions throughout the project. BJS was supported by a Henry Dale Fellowship jointly funded by the Wellcome Trust and the Royal Society (109408/Z/15/Z), and a Wellcome Trust Discovery Award (225360_Z_22_Z). DAT was funded by a BBSRC New Investigator Grant (BB/X000907/1). MJH was supported by a Leverhulme Project Grant (RPG-2018-356) and a BBSRC DTP studentship (BB/T008695/1, project reference 2599454 awarded to DAT).

## Author Contributions

MJH performed all the experiments, microscopy, and data analysis, and interpreted the data. TF contributed to the experimental design and provided technical expertise for HCR (implementation, imaging, and analysis). BJS conceived the project, obtained funding, and interpreted the data. DAT interpreted the data and wrote the paper. All authors commented on and approved the manuscript prior to submission.

---

**Figure S1:**
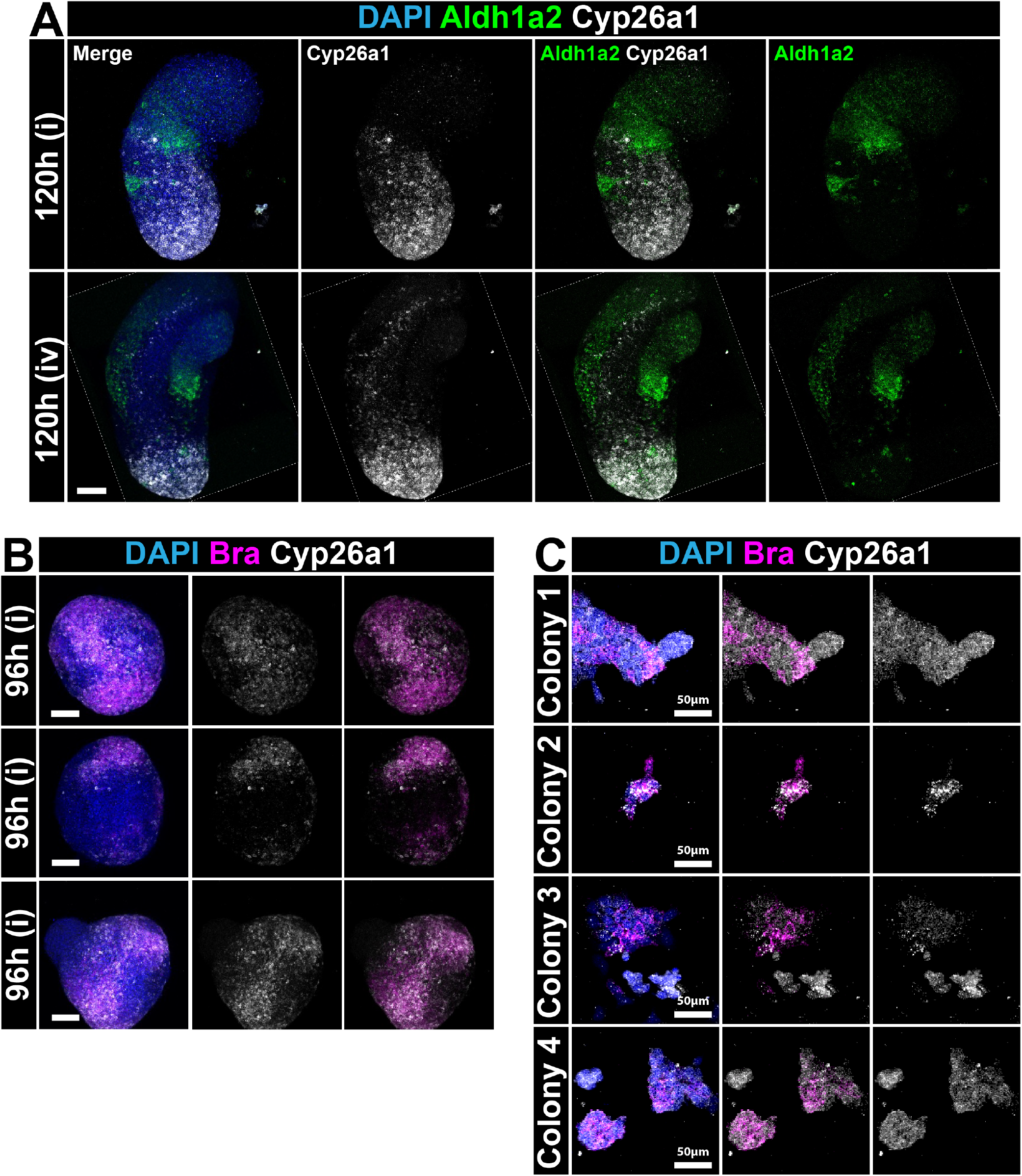
*in situ* Hybridisation Chain Reaction (HCR) analysis of *Aldh1a2, Cyp26a1*, and *Bra* in gastruloids and 2D culture. (**A**) Gastruloids from wild-type E14-Tg2A mouse ESCs as in **Fig. 1A** with the individual channels shown. (**B**) Additional examples of the expression pattern of *Bra* and *Cyp26a1* in 72h AA gastruloids. (**C**) Four locations within monolayer culture of E14-Tg2A mouse ESCs treated with continuous 3µM Chi and probed for *Bra* and *Cyp26a1*. DAPI marks the nuclei in all images. Scale bar indicates 100/*mu*m unless otherwise indicated.

**Figure S2:**
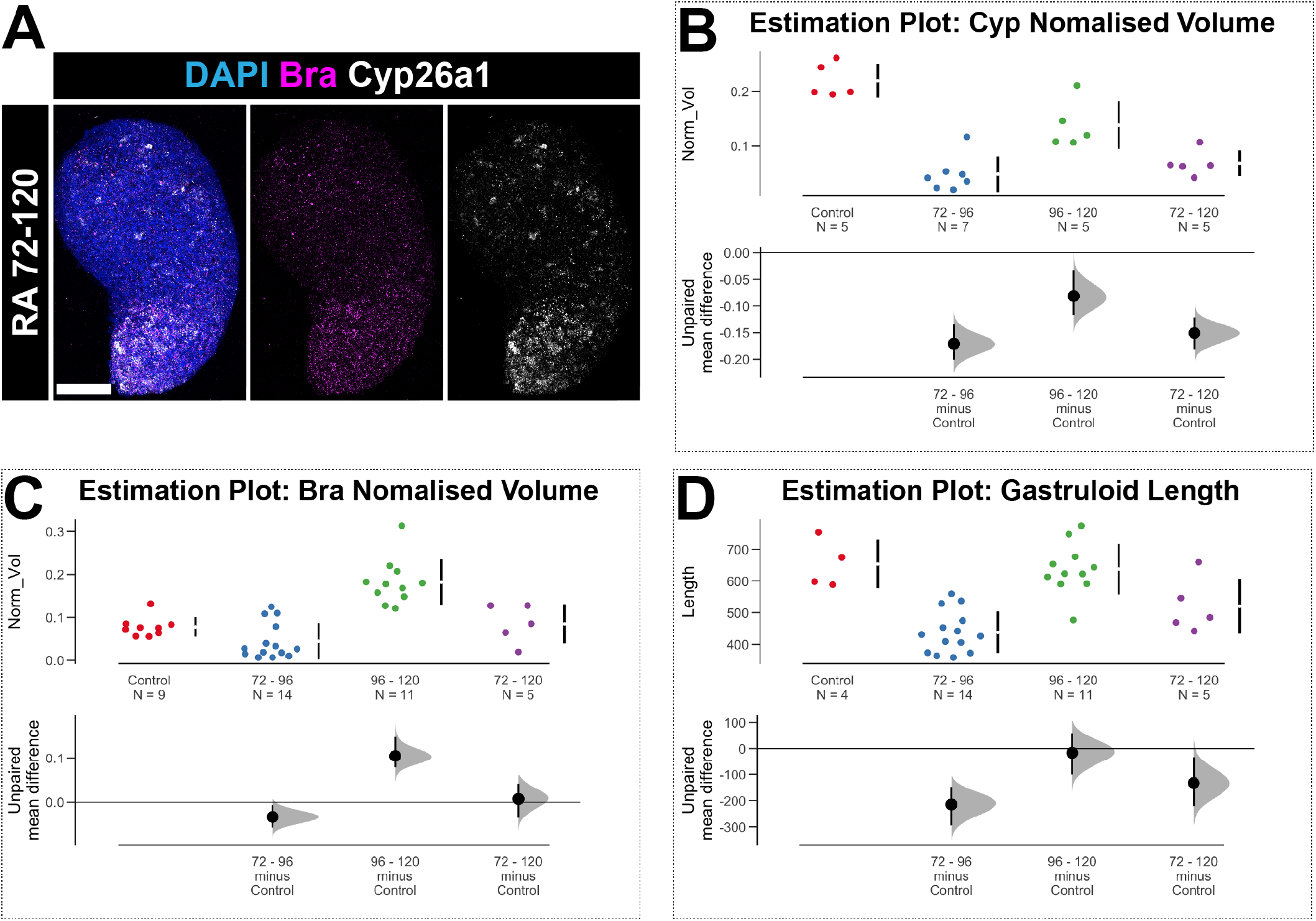
Temporal Sensitivity of gastruloids to low-dose RA. (**A**) HCR analysis of *Bra* and *Cyp26a1* following a RA pulse between 72 and 96h AA. Scale bar indicates 100µm. (**B-D**) The mean difference for three comparisons (RA pulsing between 72-96, 90-120, and 72-120h AA) against the shared control are shown in the Cumming estimation plot. The raw data for *Cyp26a1* (**B**) and *Bra* (**C**) normalised volumes, and the length of gastruloids in each treatment (**C**)are plotted on the upper axes. On the lower axes, mean differences are plotted as bootstrap sampling distributions. Each mean difference is depicted as a dot. Each 95% confidence interval is indicated by the ends of the vertical error bars.

**Figure S3:**
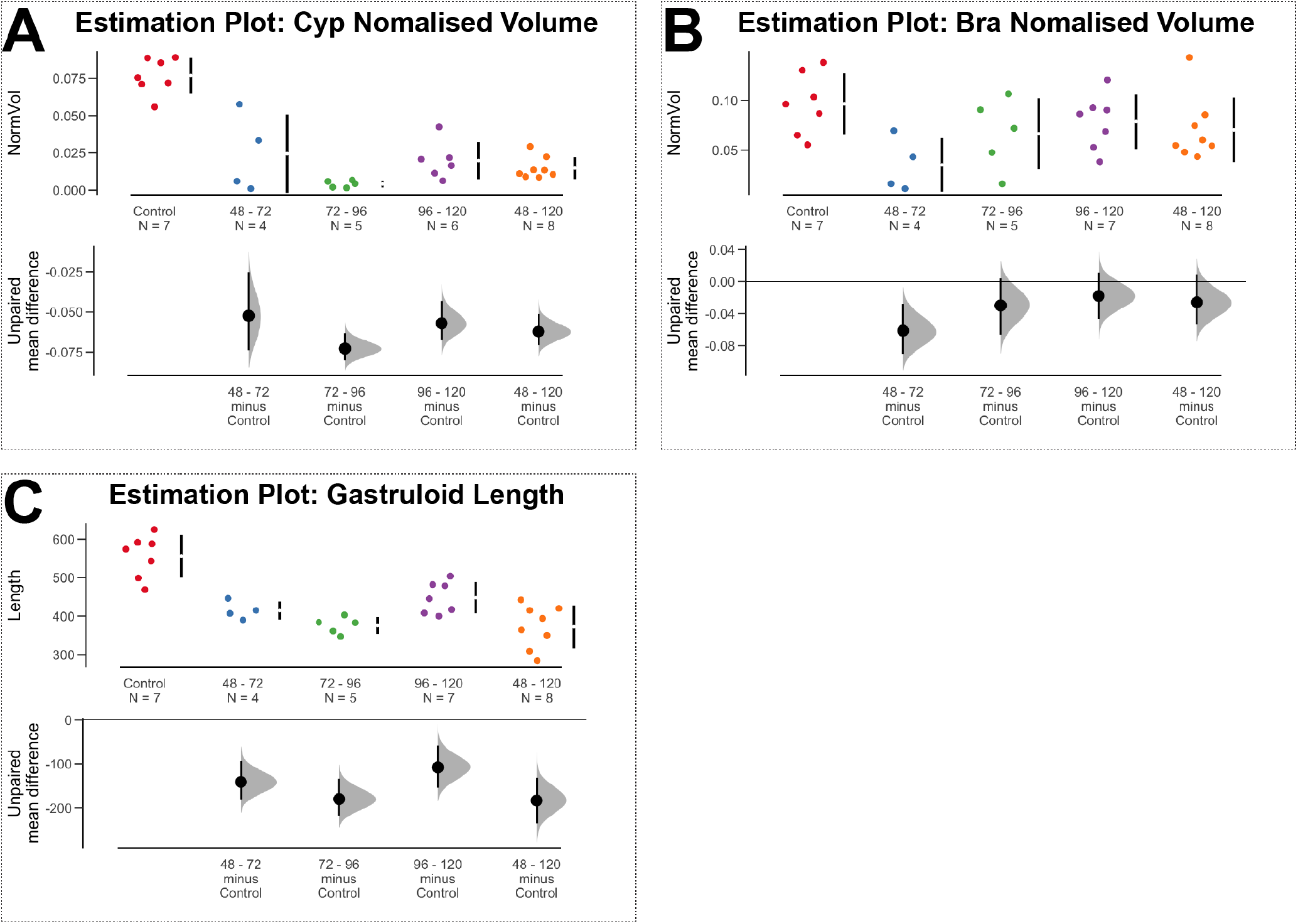
Cumming Estimation Plots for gastruloids treated with AGN. Cumming estimations plots analysing the normalised volumes of *Cyp26a1* (**A**) and *Bra* (B), as well as the lengths of gastruloids (**C**) following durations of the Retinoic Acid Receptor Inhibitor AGN. The mean difference for four comparisons (AGN pulsing between 48-72, 72-96, 96-120, 48-120h AA) against the shared control are shown. The raw data for *Cyp26a1* (**B**) and *Bra* (**C**) normalised volumes, and the length of gastruloids in each treatment (**C**)are plotted on the upper axes. On the lower axes, mean differences are plotted as bootstrap sampling distributions. Each mean difference is depicted as a dot. Each 95% confidence interval is indicated by the ends of the vertical error bars.

**Table S1:**
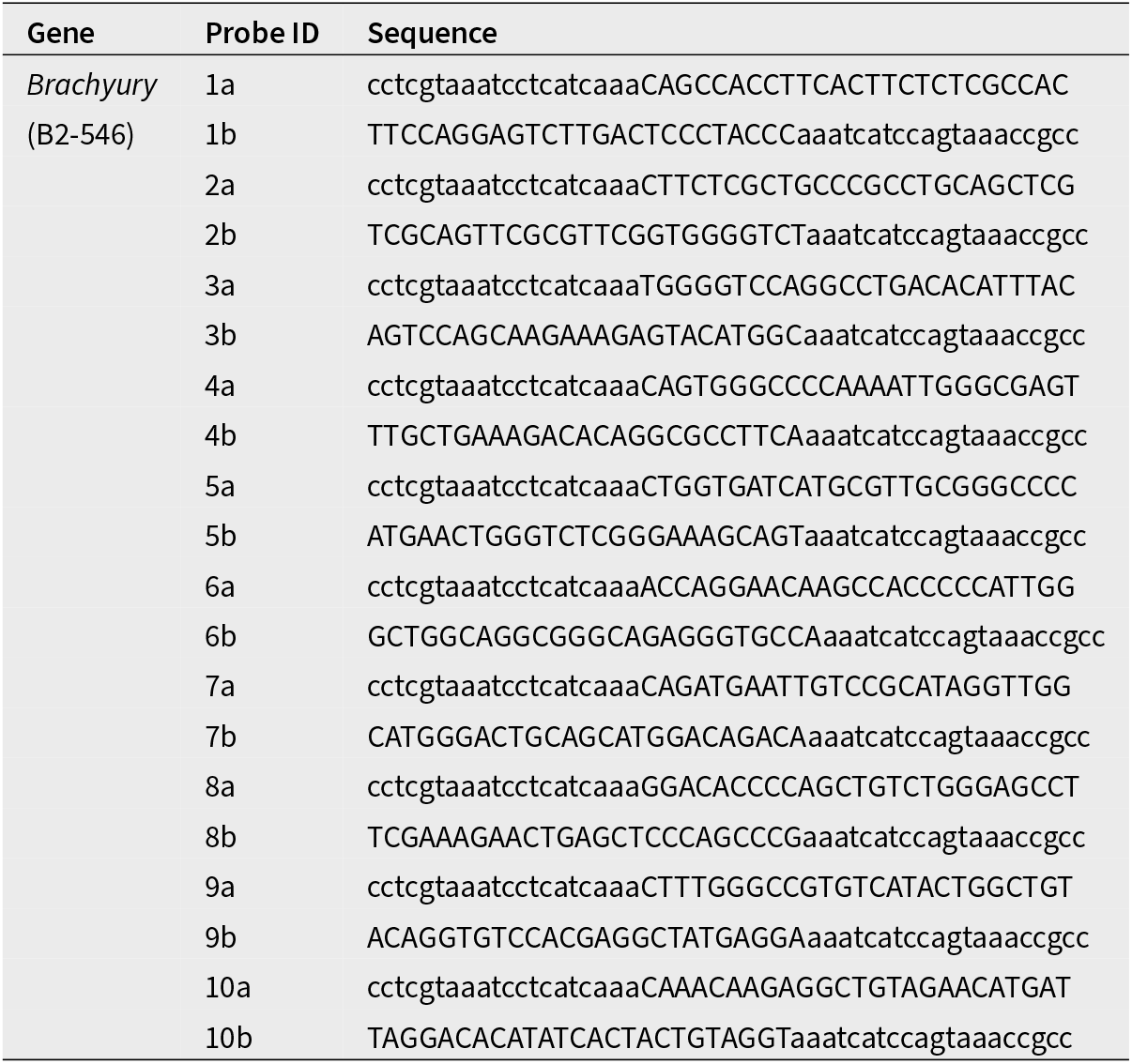
In Situ Hybridisation Chain Reaction probes and hairpins for *Brachyury*.

**Table S2:**
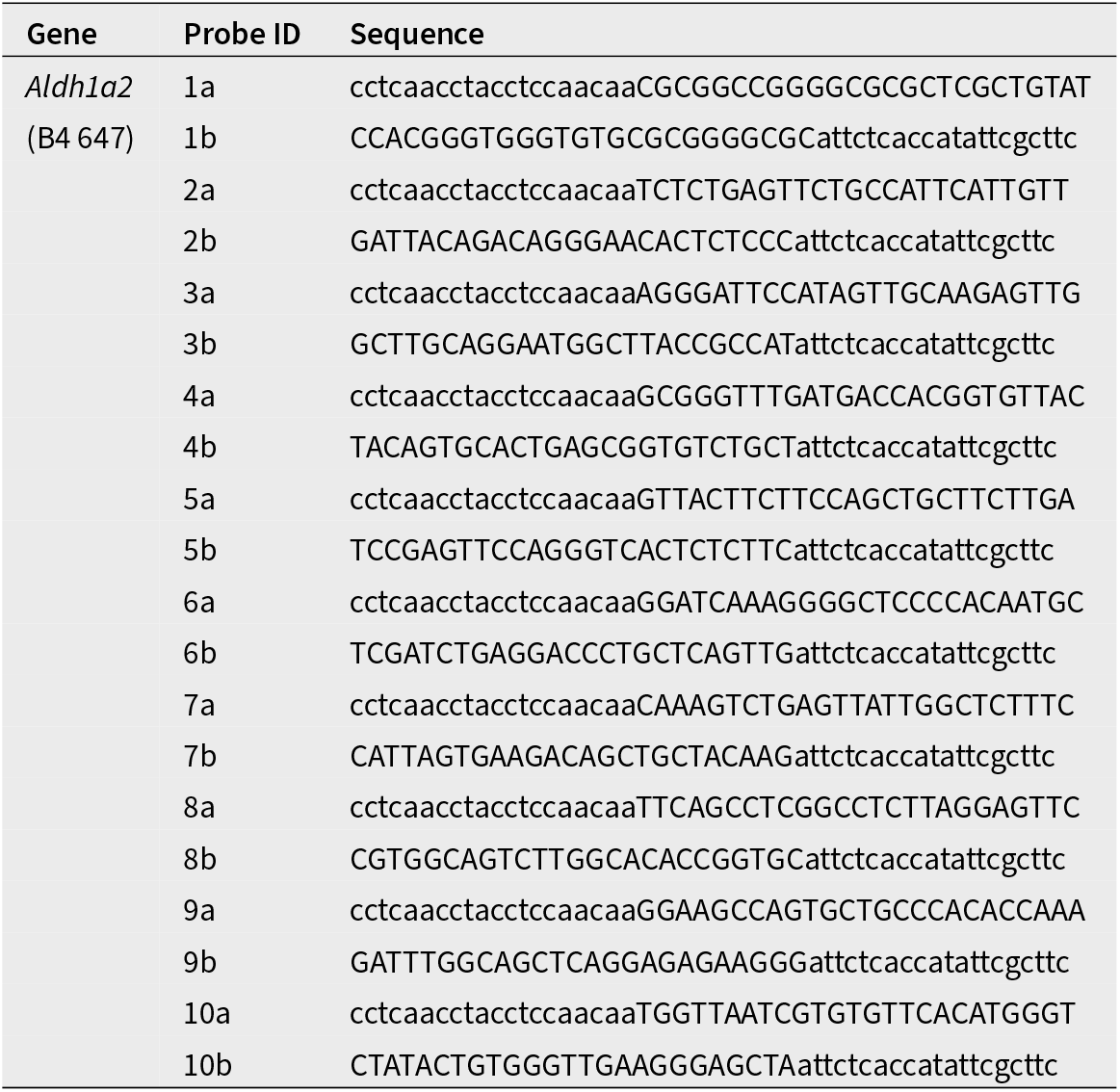
In Situ Hybridisation Chain Reaction probes and hairpins for *Aldh1a2*.

**Table S3:**
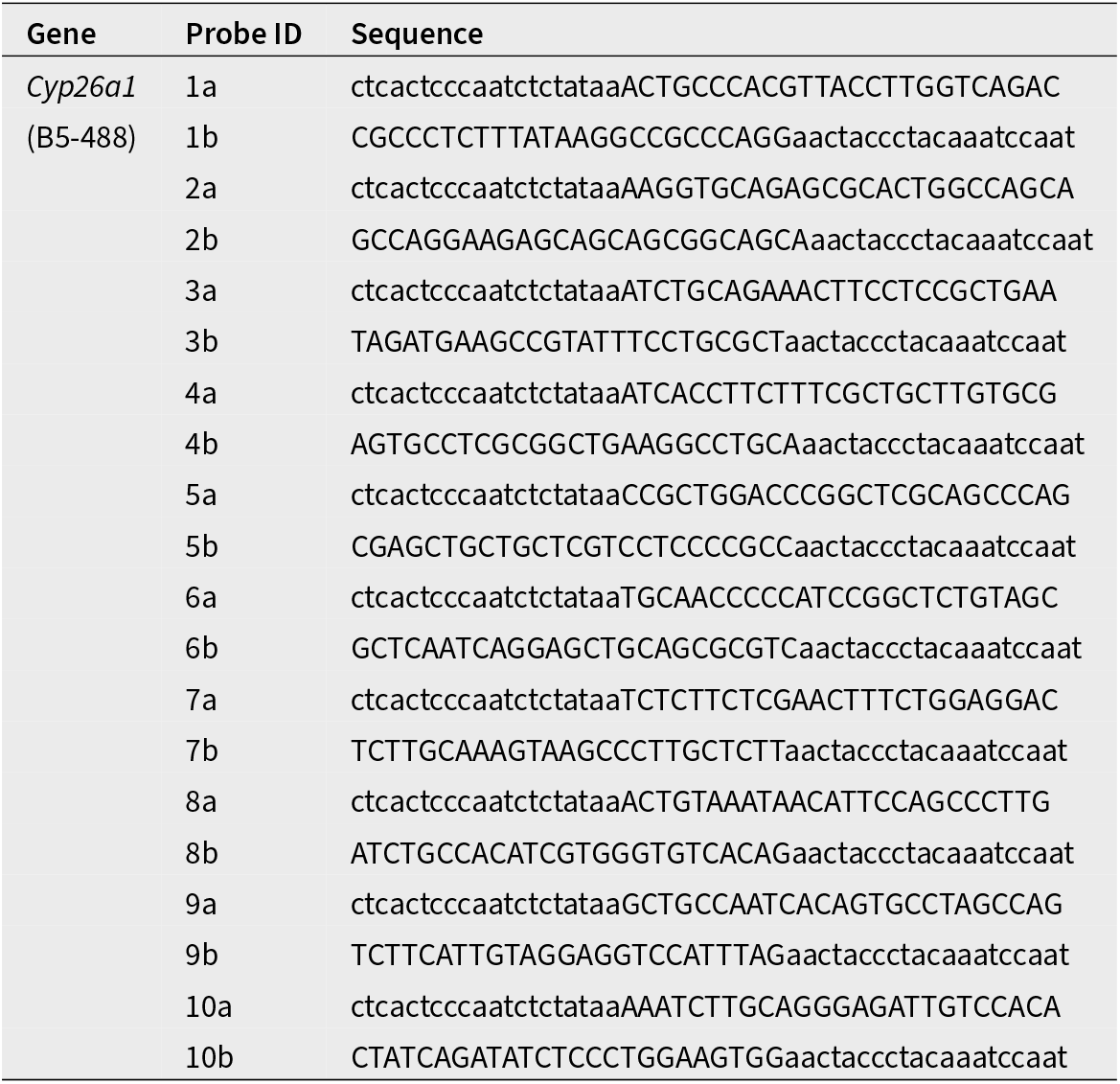
In Situ Hybridisation Chain Reaction probes and hairpins for *Cyp26a1*.

